## Supplementary figures and images for "Practical Fluorescence Reconstruction Microscopy for Large Samples and Low-Magnification Imaging"

### Supplemental Figure 1

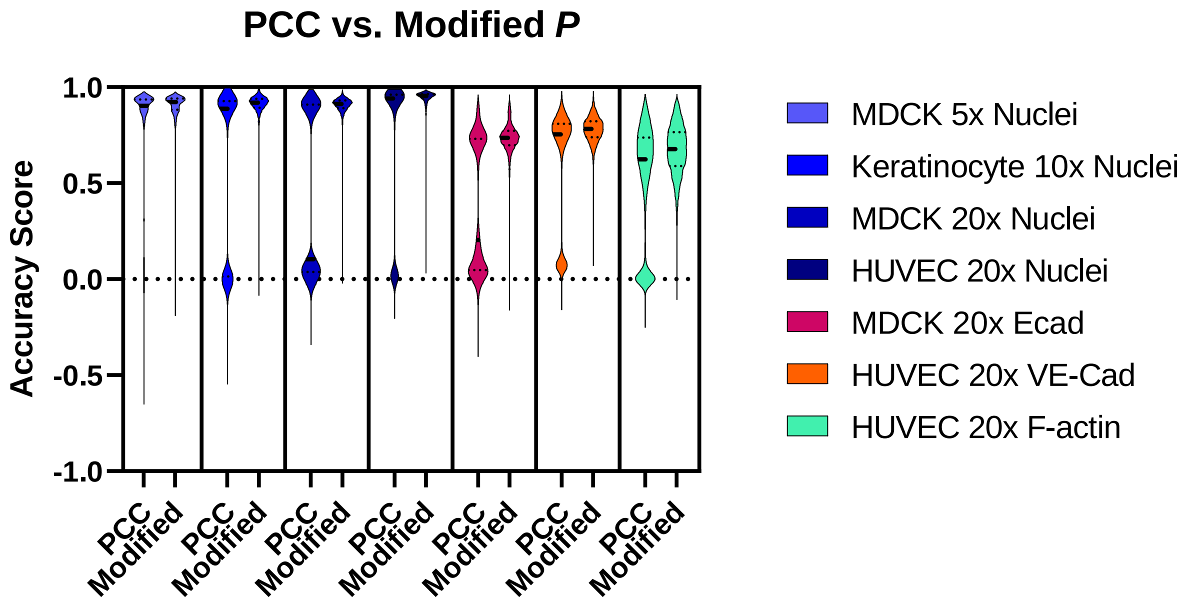

### Supplemental Figure 2

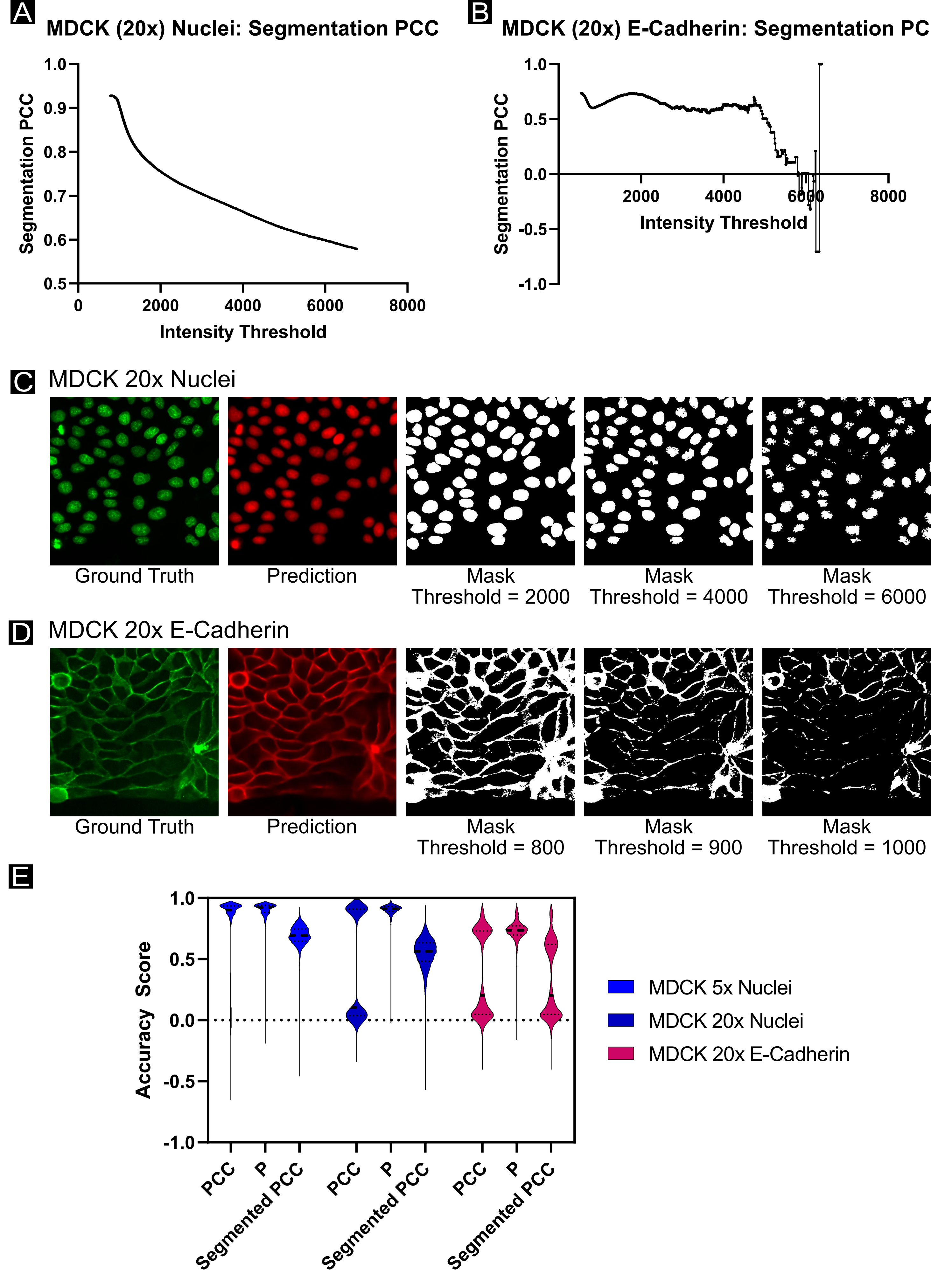

### Supplemental Figure 3

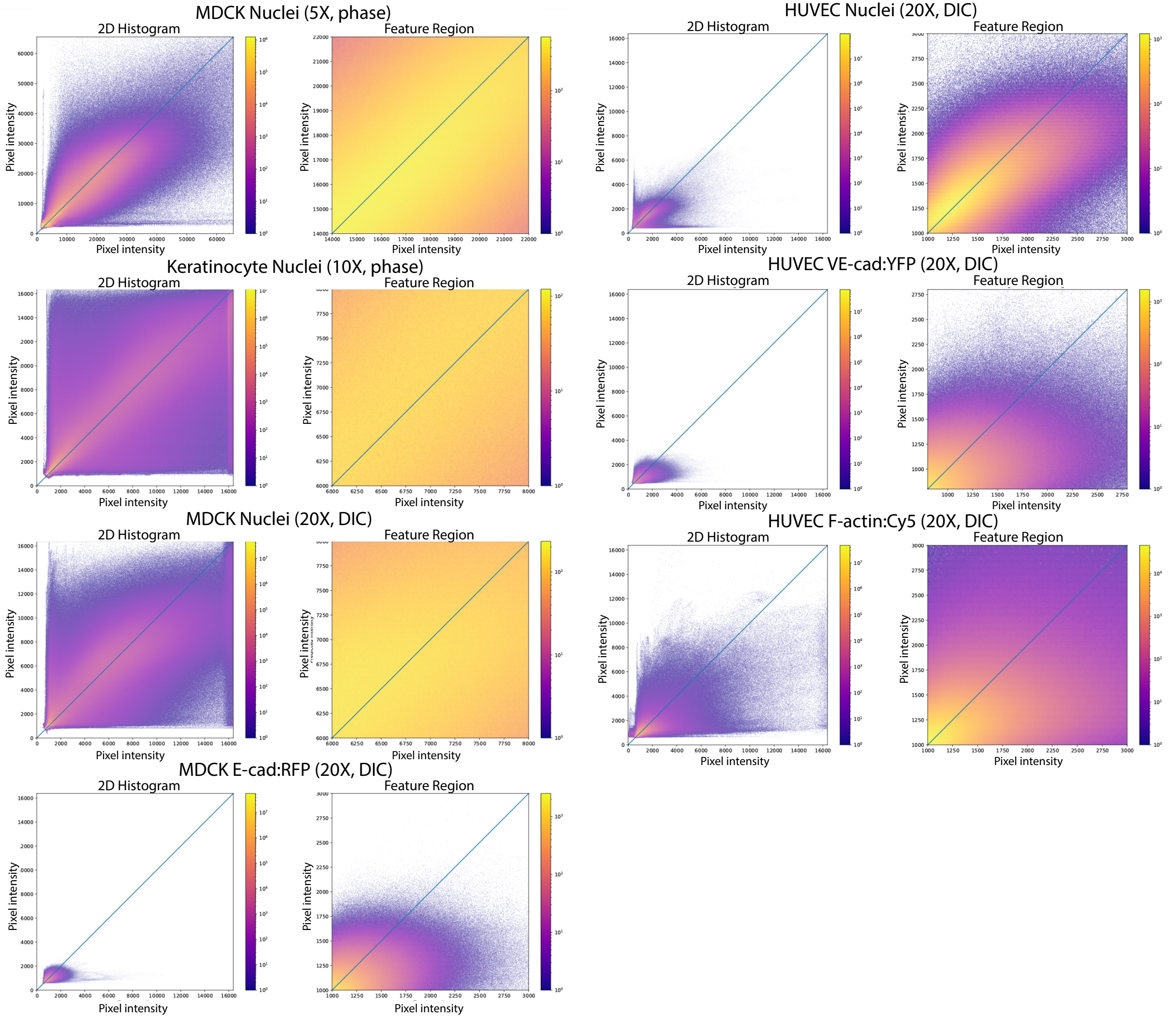

### Supplemental Figure 4

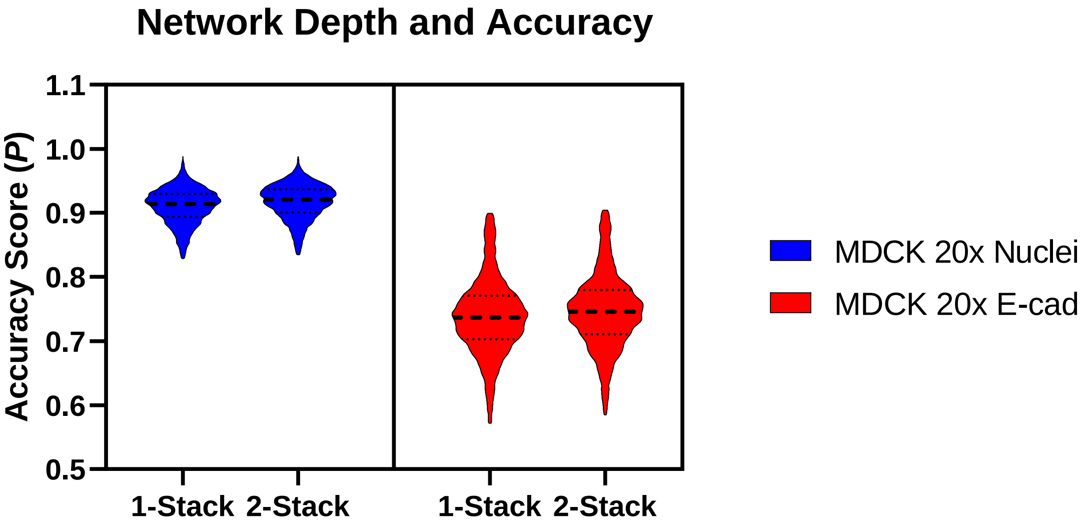

### Supplemental Figure 5

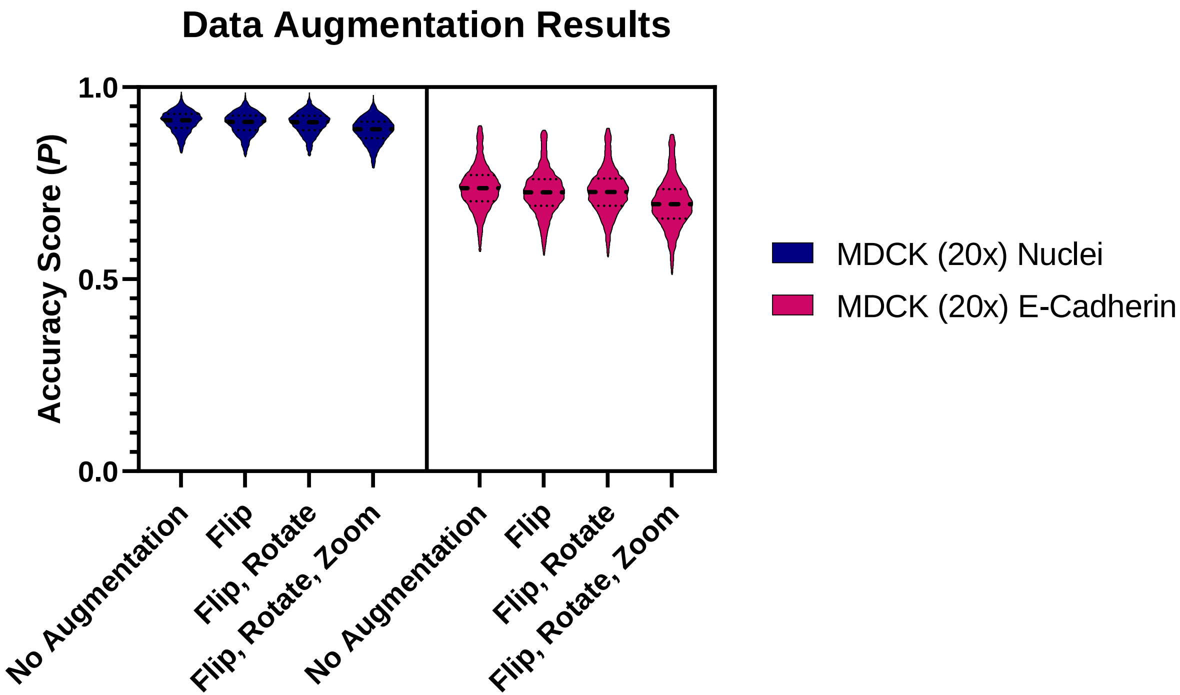

### Supplemental Figure 6

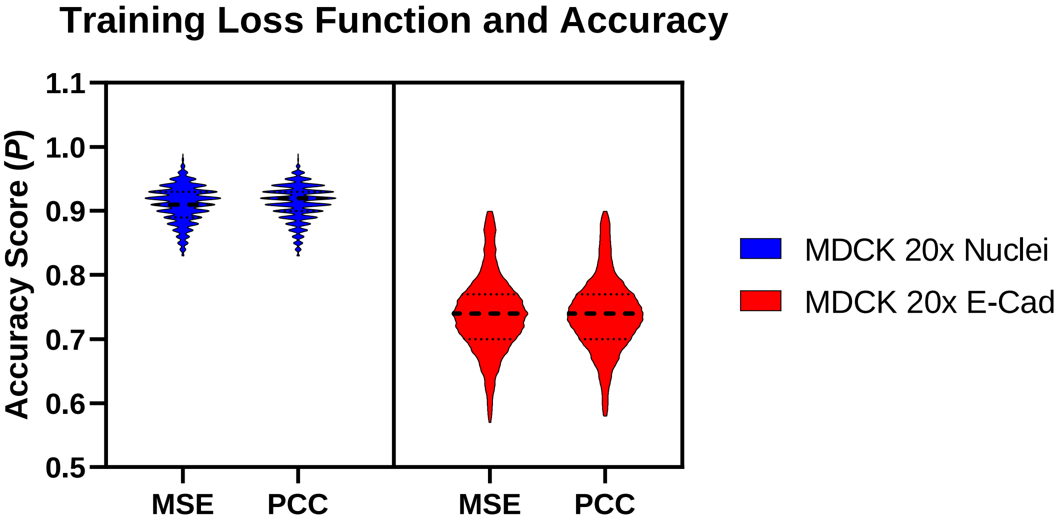

### Supplemental Figure 7

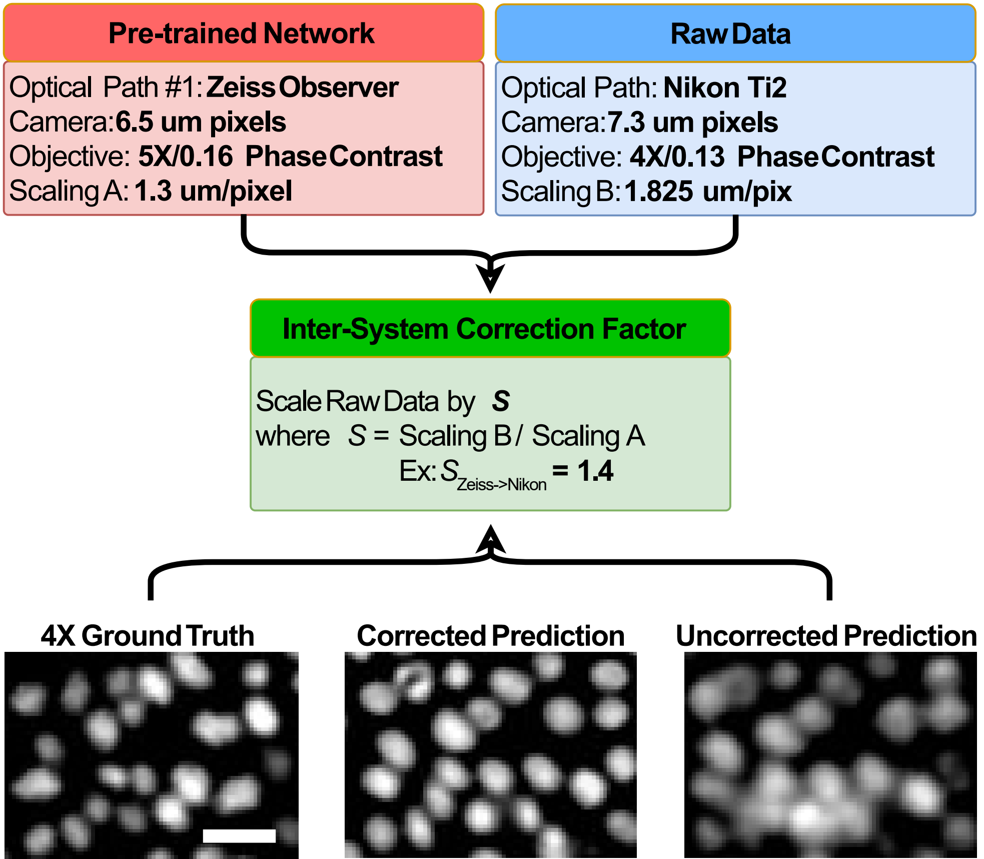

### Supplemental Figure 8

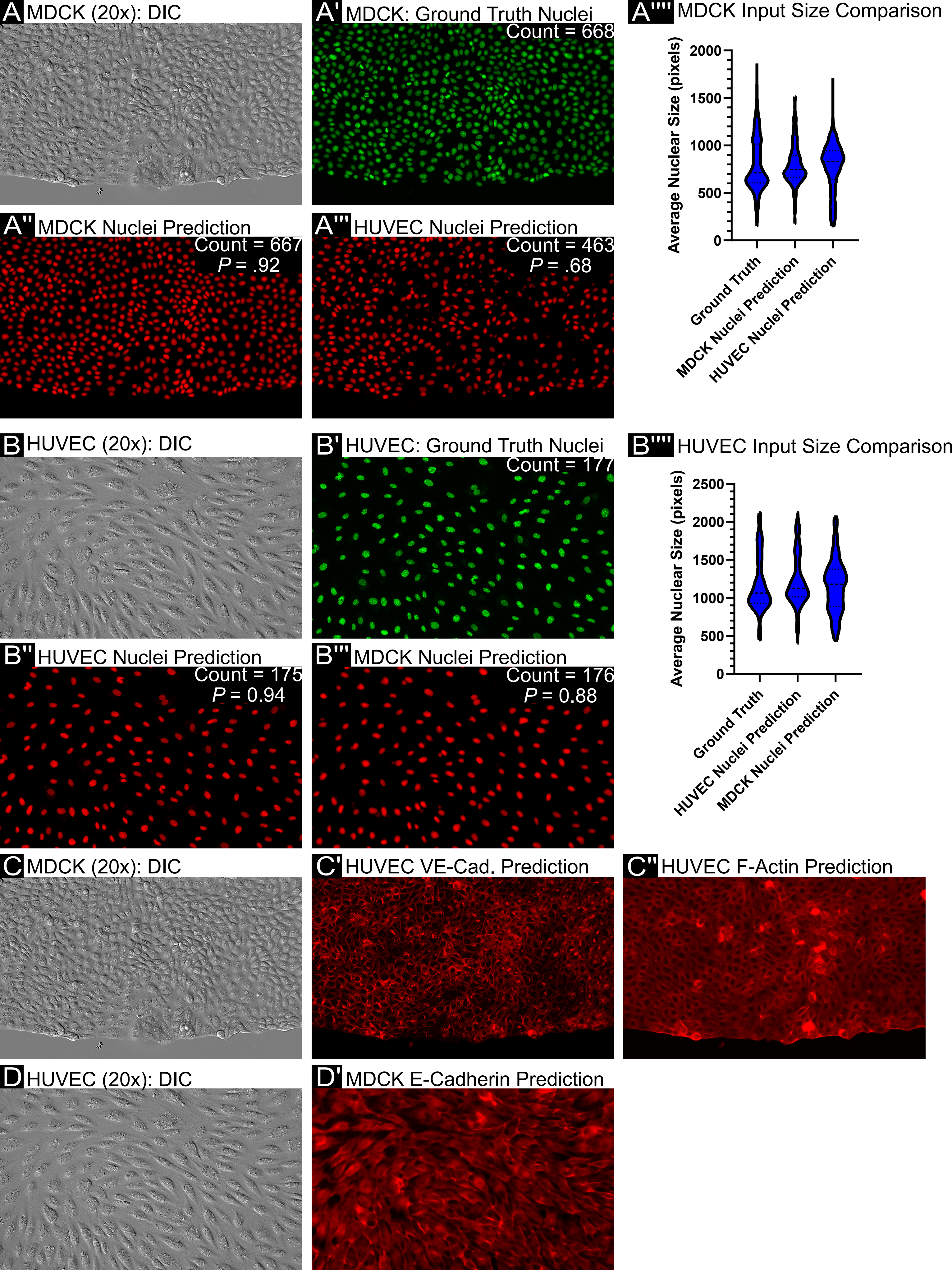

### Supplemental Figure 9

#### Training and Validation Loss: Keratinocyte 10x Nuclei

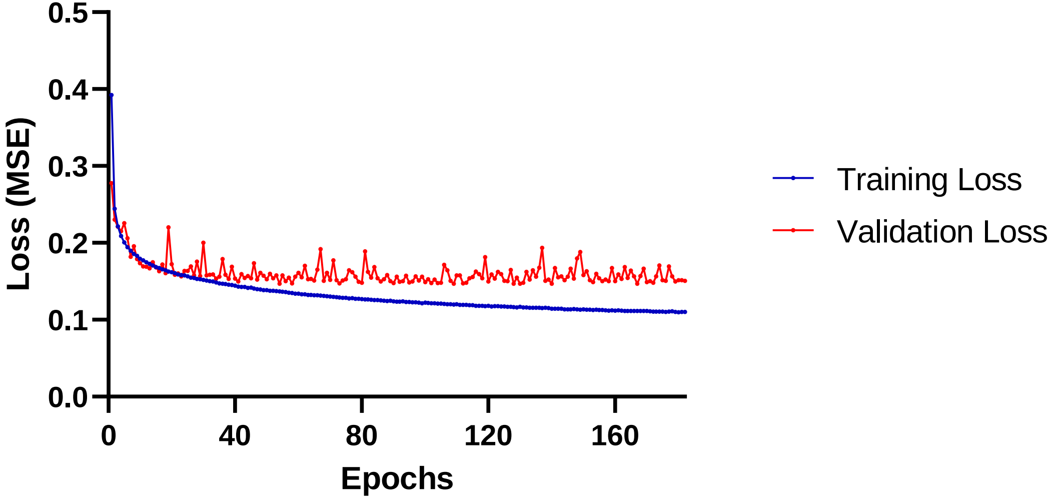
